# Dose-dependent role of Gfi1 in murine hematopoietic stem cell self-renewal and differentiation

**DOI:** 10.1101/625715

**Authors:** Judith Schütte, Aniththa Thivakaran, Yahya Al-Matary, Pradeep Kumar Patnana, Daria Frank, Daniel R. Engel, Ulrich Dührsen, Cyrus Khandanpour

## Abstract

Gfi1 (Growth factor independence 1) is a transcription factor that influences the stem cell capacity of hematopoietic stem cells (HSCs) as well as their differentiation into the myeloid and lymphoid lineage. Loss of Gfi1 impedes the repopulation capacity of HSCs and leads to a block in granulocyte generation causing severe neutropenia and monocytosis. Competitive transplantation assays showed that Gfi1-deficient cells were not able to reconstitute myeloid and lymphoid hematopoiesis in competition with Gfi1-wildtype (*GFI1*-36S) cells. Low Gfi1 levels (*GFI1*-knockdown = *GFI1*-KD) in blasts of myelodysplastic neoplasms, acute and chronic myeloid leukemia patients are associated with poor patient survival. To understand how reduced levels or loss of Gfi1 contribute to hematopoiesis, we analyzed the effect of *GFI1*-KD and *Gfi1*-KO on HSCs and more mature cell types in mice. *GFI1*-KD and *Gfi1*-KO led to strong decrease in HSC numbers, while the numbers of early progenitors (Lin^−^ Sca1^+^cKit^+^ cells) were slightly increased. Competitive transplantation assays showed that *GFI1*-KD and *Gfi1*-KO HSCs can still engraft and expand, but they cannot contribute to myeloid and lymphoid differentiation.

## Introduction

Gfi1 (Growth factor independence 1) is a transcriptional repressor with a crucial role in hematopoietic stem cells (HSCs) as well as more mature cells of the myeloid and lymphoid lineage (1, 2). It regulates maturation of myeloid cells and is required for different signaling pathways such as Notch signaling (3–5). Analysis of *Gfi1* expression in *Gfi1*:GFP heterozygous knock-in mice revealed that *Gfi1* is primarily expressed in HSCs, GMPs (granulocyte-macrophage progenitors) and CLPs (common lymphoid progenitors), while it is absent in CMPs (common myeloid progenitors) and MEPs (megakaryocyte-erythroid progenitors) (6). Gfi1-deficient mice are characterized by an accumulation of monocytic cells and absence of granulocytes, leading to severe neutropenia and monocytosis (7, 8). Recently it has been implicated that reduced levels of Gfi1 are associated with an inferior prognosis for human MPN (myeloproliferative neoplasm), CML (chronic myeloid leukemia) and AML (acute myeloid leukemia) patients and with an acceleration of AML development in mice (9–11). These studies also showed that Gfi1 regulates expression of certain oncogenes, apoptotic pathways and possibly metabolic functions in a dose-dependent manner. However, it has also been postulated that loss of Gfi1 negatively influences the repopulation capacity of HSCs (6, 12).

We were now interested whether low levels and loss of Gfi1 indeed negatively affect the stem cell and differentiation capacity of HSCs and how the seemingly contradictory findings between reduced self-renewal capacity and the dose-dependent role of Gfi1 function in myeloid pathogenesis could be reconciled. For the analysis we made use of *Gfi1*-knockout (KO) mice, a mouse strain with a complete loss of Gfi1, and *GFI1*-knockdown (KD) mice, a mouse strain in which we cloned the human *GFI1* sequence into the murine *Gfi1* gene locus together with a Neo cassette in the opposite direction of transcription, leading to *GFI1* expression of only 10-15% of wild-type levels (10). We show here that *GFI1*-KD and *Gfi1*-KO mice contain a higher number of hematopoietic progenitors (LSK cells, Lin^−^Sca1^+^c-Kit^+^), but HSC (Lin^−^Sca1^+^c-Kit^+^CD150^+^CD48^−^) numbers are strongly reduced. Although *GFI1*-KD and *Gfi1*-KO mice have fewer HSCs compared to *Gfi1*-WT mice, these HSCs can still engraft and expand in a competitive setting. However, they have lost their ability to contribute to multi-lineage differentiation.

## Methods

### Mice

*GFI1*-36S, *Gfi1*-KO and *GFI1*-KD mice have previously been described (8, 10, 13). Mice were housed in specific pathogen-free conditions at the animal facility of University Hospital Essen, Germany. All experiments were approved by the local authorities (LANUV, Az84-02.04.2015.A022).

Mice were irradiated using the X-Rad320 device from Precision X-ray. GFP-sorted whole BM cells or Lineage negative cells were transplanted intravenously.

### Lineage negative cell isolation

BM cells were isolated from tibiae, femora and humeri and red blood cells were removed by using BD Pharm Lyse™ (555899, BD Biosciences). Lineage negative cells were collected by applying the Direct Lineage Cell Depletion Kit mouse (130-090-858, Miltenyi Biotec GmbH) and magnetic separation following manufacturer’s instructions.

### Flow Cytometry

The following antibodies were used to differentiate the various cell types: Lineage Cell Detection Cocktail-Biotin, mouse (130-092-613) from Miltenyi Biotec GmbH, FITC anti-mouse Ly-6A/E (Sca-1) (108105), Brilliant Violet 421™ anti-mouse CD117 (c-Kit) (105827), PerCP/Cy5.5 anti-mouse CD45.1 (110728), PE anti-mouse CD45.2 (109808), PE/Cy7 anti-mouse CD150 (SLAM) (115913), APC anti-mouse CD48 (103411), APC anti-mouse/human CD11b (101211), APC anti-mouse CD8a (100711), FITC anti-mouse CD4 (100509) and FITC anti-mouse/human CD45R/B220 (103205) from BioLegend, Streptavidin PerCP-Cyanine5.5 (45-4317-82) and Ly-6G/Ly-6C Monoclonal Antibody (RB6-8C5) FITC from eBioscience. Cells were analysed using a BD LSR II device from BD Biosciences.

### Statistical Analysis

Analyses were performed using Microsoft Excel. Significance was calculated using a paired t-test with a normal distribution.

All methods were performed in accordance with the relevant guidelines and regulations.

## Results and Discussion

GFI1 influences prognosis of AML, CML and MPN patients in a dose-dependent manner, and in murine models of human leukemia reduced levels of GFI1 accelerated disease development (9–11, 14). AML, CML and MPN arise as a result of transformation of HSCs or early hematopoietic progenitors that still possess some degree of stem cell capacity (15). It has been described that complete loss of Gfi1 leads to an impaired function of HSCs (6, 7) and an increase in LSK frequency, possibly due to an increased proliferation (7, 16). It has previously been reported that *Gfi1*-KO cells fail to compete with *Gfi1*-WT cells in a competitive setting, demonstrated by a lack of reconstitution capacity of the myeloid and lymphoid compartment in the peripheral blood (PB) (6). The question arising from these results is whether the lack of mature cells in a competitive transplantation setting is a) due to a failure of *Gfi1*-KO HSCs to engraft, b) due to a lack of survival of *Gfi1*-KO HSCs in competition with *Gfi1*-WT HSCs or c) due to a loss of *Gfi1*-KO HSC capacity to contribute to PB cell production. We were hence interested to determine whether reduced levels or loss of GFI1/Gfi1 influence a) the number of HSCs and b) their ability to engraft and reconstitute the PB and BM of recipient mice in a competitive transplantation setting.

To this end, we made use of the previously described *GFI1*-KD mice which express human GFI1 at around 10-15 % of normal GFI1 levels (10) and *Gfi1*-KO mice with a complete loss of Gfi1 expression (8). As a control we used *GFI1-36S* mice, a mouse strain containing the human *GFI1* cDNA sequence at the position of the murine *Gfi1* locus, as this is the mouse strain from which *GFI1*-KD mice were derived. We have shown in the past that *GFI1*-36S mice are functionally equivalent to murine *Gfi1-WT* mice (10, 13, 17, 18). In addition, they do not differ in their number of hematopoietic stem and progenitor cells (Supplementary Figure 1 A and B). We first examined the frequency of hematopoietic progenitor cells in *GFI1*-36S, *GFI1*-KD and *Gfi1*-KO mice. While the percentage of Lin^−^Sca1^+^c-Kit^+^ (LSK) cells was significantly increased in *GFI1*-KD mice compared to *GFI1-36S* mice, the number of LSKs in *Gfi1*-KO mice was only marginally elevated (Fig. 1A and B). In contrast, the percentage of HSCs defined as Lin^−^Sca1^+^c-Kit^+^CD150^+^CD48^−^ was highly reduced in both, *GFI1*-KD and *Gfi1*-KO mice (Fig. 1 A and C). This observation was irrespective of the age of the mice (Supplementary Fig. 1 C and D). Our data thus recapitulate previous findings of our and other groups regarding an expansion of LSKs upon loss of *GFI1* (7, 16), but further expands our knowledge about a dose-dependent effect of *GFI1* expression on HSCs, namely a statistically significant, stepwise decrease in HSC numbers.

**Figure 1:**
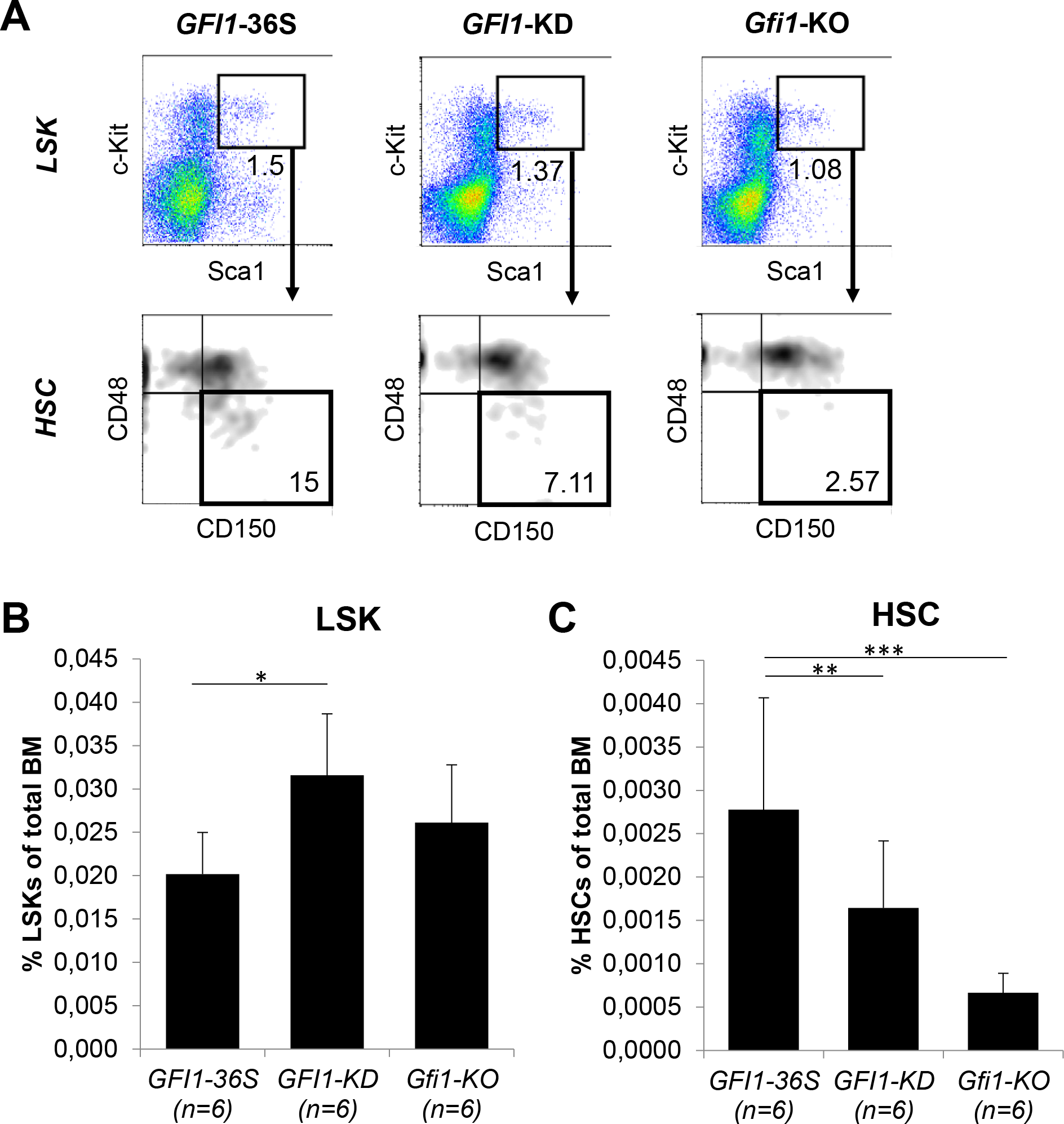
Gfi1 is required for HSC generation. **A. Gating strategy for identification of HSCs.** Murine BM cells were isolated and sorted for surface marker expression (lineage marker, c-Kit, Sca1, CD48 and CD150) by flow cytometry. Flow cytometry data for one mouse per genotype (*GFI1*-36S, *GFI1*-KD and *Gfi1*-KO) are shown. **B.** Reduced levels of GFI1 (*GFI1*-KD) and loss of Gfi1 (*Gfi1*-KO) led to a slight increase in LSK (Lin^−^Sca1^+^c-Kit^+^) cells. Depicted is the percentage of LSK cells compared to total BM cells. For each genotype six mice were analyzed. Error bars represent standard deviation. *=p-value ≤ 0.05. **C.** Knockdown or loss of GFI1/Gfi1 resulted in a dose-dependent decrease of HSCs (Lin^−^Sca1^+^c-Kit^+^CD150^+^CD48^−^). Cell isolation and analysis were performed as described in A. Shown is the percentage of HSCs compared to total BM cells. For each genotype six mice were analyzed. Error bars represent standard deviation. * ≤ 0.05, **=p-value ≤ 0.01, ***=p-value ≤ 0.001

We next investigated how reduced levels or loss of GFI1 in HSCs affect their ability to engraft after transplantation and to differentiate into the various cell lineages. To study this we performed a competitive bone marrow (BM) transplantation assay (Fig. 2A). We isolated Lin^−^ cells from CD45.1-expressing *Gfi1*-WT mice and mixed those cells in equal numbers with Lin^−^ cells from CD45.2-expressing *GFI1*-KD or *Gfi1*-KO mice. Next, we transplanted a total of 10^5^ Lin^−^ cells into lethally irradiated C57BL/6 mice and analysed the PB and BM for their CD45.1 and CD45.2 expression. Although *GFI1*-KD and *Gfi1*-KO mice contain fewer HSCs than *Gfi1*-WT mice, we transplanted equal numbers of Lin^−^ cells, the cell fraction that contains hematopoietic stem and progenitor cells. We chose to transplant Lin^−^ cells and not phenotypically more defined HSCs because it has recently been shown that altered transcription factor levels can influence which cell fractions give rise to long-term repopulation. The combined deletion of *p16*^*Ink4a*^, *p19*^*Arf*^ and *Trp53* in mice for instance led to the acquisition of stem-cell-like properties by multipotent progenitors (19). Repeating the transplantation experiments with highly purified HSCs was not allowed due to animal regulations demanding a reduction of mouse numbers. Transplantation of Lin^−^ cells thus results in a potential bias towards transplantation of fewer *GFI1*-KD or *Gfi1*-KO HSCs (based on surface marker expression) compared to *Gfi1*-WT HSCs, but ensures that all functional cells are included in the transplanted cell fraction. To determine the contribution of *Gfi1*-WT or *GFI1*-KD/*Gfi1*-KO HSCs to hematopoiesis, we analyzed not only the overall presence of CD45.2^+^ cells in PB, but also the presence of short-lived CD45.2^+^ myeloid cells such as monocytes over time. The presence of monocytes derived from CD45.2^+^ *GFI1*-KD or *Gfi1*-KO cells in PB more than 6 weeks after transplantation rules out a potential carry-over of these cells as a result of the transplantation. To check whether *GFI1*-KD and *Gfi1*-KO derived cells possess HSC activity, we determined the presence of short-lived monocytes in PB. About 16%, 12% or 5% of all monocytes were CD45.2^+^ 77, 97 or 132 days after transplantation, respectively, hence originated from *GFI1*-KD HSCs in the competitive transplantation assay (Fig. 2 B). As it has previously been shown that long-term reconstitution can be determined 12-16 weeks after transplantation (19, 20), this result indicates that *GFI1*-KD HSCs possess indeed long-term HSC activity and that GFI1 has a dose-dependent effect in contribution to peripheral blood cell formation. In contrast, CD45.2-derived monocytes were not present in the *Gfi1*-KO transplantation experiment, even after 21 days of transplantation (Fig. 2 C). This finding is expected as it has been shown in previous experiments that *Gfi1*-KO cells do not contribute to mature cells in PB in a competitive transplantation setting (6, 12). In addition to monocytes, we also measured the overall contribution of CD45.2 transplanted cells to the PB composition. On day 308 after transplantation of *GFI1*-KD cells together with *Gfi1*-WT cells, almost 90 % of all cells in the PB had a CD45.1 origin (Fig. 2B). A very similar pattern could be observed 112 days after transplantation of *Gfi1*-KO cells together with *Gfi1*-WT cells (Fig. 2C). Once CD45.2-derived short-lived monocytes were not detected anymore in PB (after 308 days for *GFI1*-KD and after 112 days for *Gfi1*-KO), we stopped the experiment and analyzed the BM for presence of HSCs. Interestingly, two out of four transplanted mice that were generated from CD45.2-expressing *GFI1*-KD cells contained a high percentage of HSCs in the BM (44 and 88 % respectively), while the HSCs of the other two mice originated from CD45.1-expressing *Gfi1*-WT cells (Fig. 2D). Similarly, 25-40% of HSCs arose from CD45.2-expressing cells in three out of four mice transplanted with equal numbers of *Gfi1*-KO and *Gfi1*-WT cells, thus arising from *Gfi1*-KO cells (Fig. 2E). In contrast to the origin of HSCs, most of the mature cell types of the PB developed from CD45.1-expressing *Gfi1*-WT cells at 308 days in *GFI1*-KD or 112 days in *Gfi1*-KO competitive transplantation experiments (Fig. 2F and G). At day 308 after transplantation, 0.3% of the Ly6G^hi^CD11b^+^ (granulocytes) and 1.2% of Ly6G^int^CD11b^+^ (monocytes) originated from *GFI1*-KD HSCs. This lack of contribution of HSCs to more committed cells was already observable at an early timepoint of the hematopoietic hierarchy as the majority of LSKs in both competitive transplantation experiments were CD45.2-negative (Supplementary Fig. 2). We also examined the contribution of HSC to other compartments. After 308 days, CD45.2-expressing *GFI1*-KD cells still contributed to 4 % of all CD4^+^ cells, 13.5 % of all CD8^+^ cells and 7.5 % of all B220^+^ cells in PB (Fig. 2F). While B220^+^ cells only very rarely arised from CD45.2-expressing *Gfi1*-KO cells, these cells still contributed to 12 % of all CD4^+^ and 23 % of all CD8^+^ cells 112 days after transplantation (Fig. 2G). It is more difficult to estimate the contribution of transplanted HSCs to the cells of the lymphoid lineage, as these cell types are generally long-lived. They could thus originate from the host. Alternatively, if they were generated from the transplanted HSCs early after transplantation, these cells would still be detectable at later time-points, even though HSCs do not self-renew and differentiate anymore. It can also not be ruled out that these cells arise from more committed progenitor cells which were co-transplanted with the Lin^−^ cells.

**Figure 2:**
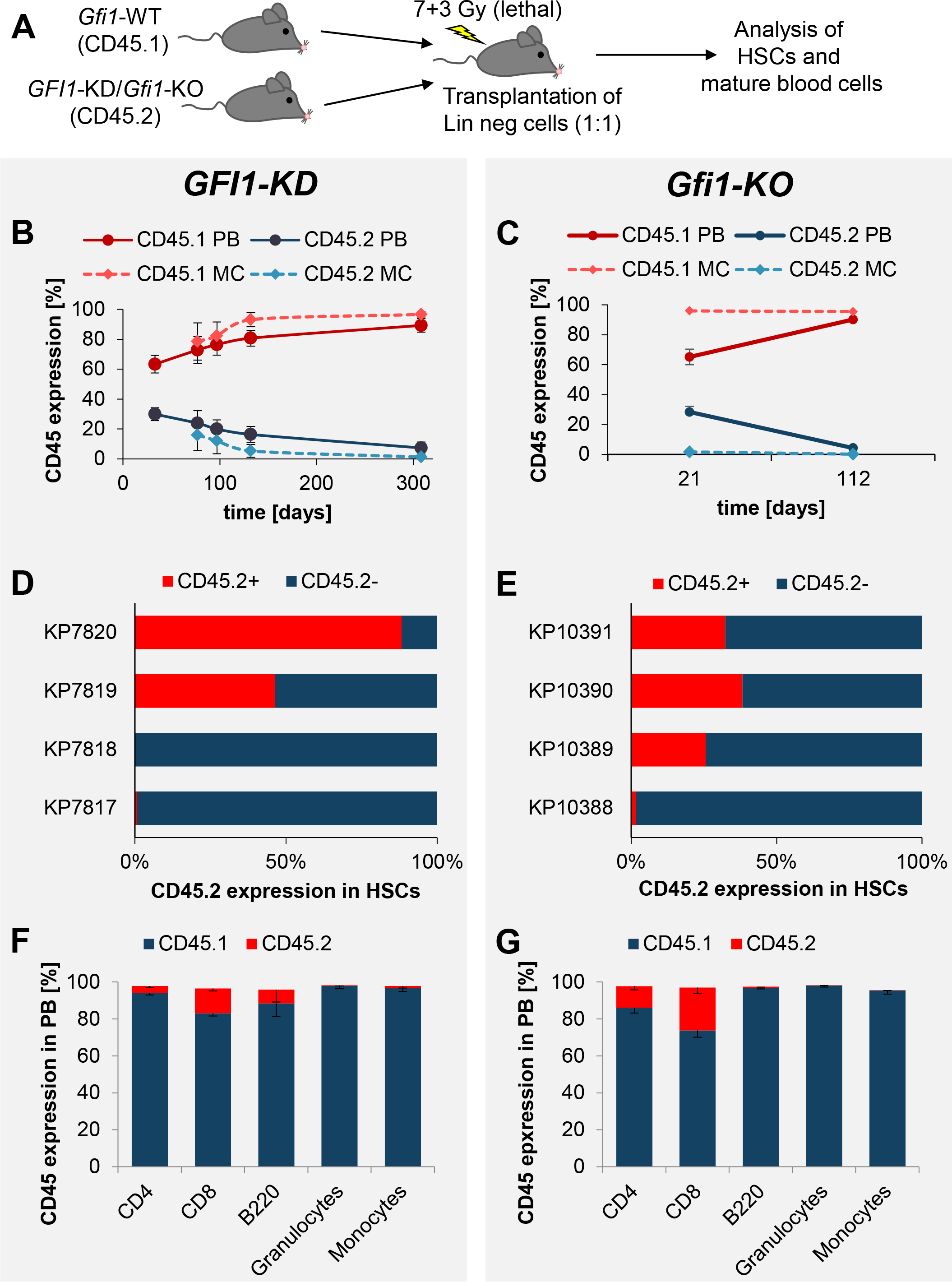
Gfi1 is essential for self-renewal of HSCs and their differentiation. **A.** Schematic diagram of the competitive transplantation assay setup. Lin^−^ cells were isolated from a CD45.1-expressing *Gfi1*-WT mouse and a CD45.2-expressing *GFI1*-KD or *Gfi1*-KO mouse. The cells were mixed in equal numbers. A total of 10^5^ Lin^−^ cells were transplanted into lethally irradiated C57BL/6 mice. For each competitive transplantation assay four mice were transplanted (n=4, 1 experiment). The BM and PB were analyzed for the presence of HSCs and mature blood cells at different time points. **B.** CD45 expression on the cell surface of all cells of the PB (PB) or more specific monocytes (MC) was measured by flow cytometry 33, 77, 97, 132 and 308 days after transplantation. **C.** CD45 expression on the cell surface of all cells of the PB or more specific monocytes (MC) was measured by flow cytometry 21 and 112 days after transplantation. **D.** CD45.2-expressing *GFI1*-KD cells contributed to the generation of HSCs (Lin^−^Sca1^+^c-Kit^+^CD150^+^CD48^−^). Four mice with the indicated names (KP7817, KP7818, KP7819 and KP7820) were subjected to transplantation and BM cells were isolated 315 days (KP7819 and KP7920) or 322 days (KP7817 and KP7818) after transplantation. The origin of HSCs was analyzed by flow cytometry. **E.** CD45.2-expressing *Gfi1*-KO cells contributed to the generation of HSCs (Lin^−^Sca1^+^c-Kit^+^CD150^+^CD48^−^). Four mice with the indicated names (KP10388, KP10389, KP10390 and KP10391) were subjected to transplantation and BM cells were isolated 112 days after transplantation. The origin of HSCs was analyzed by flow cytometry. **F.** Almost all mature blood cells of the PB had a CD45.1-expressing *Gfi1*-WT origin 308 days after transplantation of CD45.1-expressing *Gfi1*-WT and CD45.2-expressing *GFI1*-KD cells. Granulocytes defined as Ly6G^hi^CD11b^+^, Monocytes defined as Ly6G^int^CD11b^+^. Shown is the analysis of four mice. **G.** Almost all mature blood cells of the PB had a CD45.1-expressing *Gfi1*-WT origin 112 days after transplantation of CD45.1-expressing *Gfi1*-WT and CD45.2-expressing *Gfi1*-KO cells. Granulocytes defined as Ly6G^hi^CD11b^+^, Monocytes defined as Ly6G^int^CD11b^+^. Shown is the analysis of four mice. Error bars for all sub-figures represent standard deviation.

In summary, we expand our knowledge about the role of Gfi1 in HSCs by showing that reduced levels and loss of Gfi1 lead to a slight expansion of LSK cells, but to highly significant reduction of HSCs, and that this reduction is dependent on the Gfi1 level. On a functional level, reduced expression or loss of *GFI1* expression did not dramatically impede the ability of *GFI1*-KD or *Gfi1*-KO HSCs to engraft or to expand after transplantation (up to 80 % of HSCs were of *GFI1*-KD origin and up to 40% of HSCs were of *Gfi1*-KO origin), but negatively and in a dose-dependent manner influenced the maturation of HSCs into the lymphoid and myeloid lineages. Our data are in line with other reports highlighting the role of Gfi1 in lineage choice, similar to that of other transcription factors such as IRF8 and PU.1 (21, 22). Our arising hypotheses are that a) knockdown and loss of GFI1 might regulate the choice of symmetric and asymmetric division of HSCs which could explain the leukemia-propagating function of reduced GFI1 levels in CML and AML (10) or b) *GFI1*-KD or *Gfi1*-KO HSCs exhaust their ability to produce mature cells after a certain time point. As *Gfi1*-KO cells have a normal life span in a non-competitive setting in mice if they are kept under SPF conditions (23), further experiments are needed to study the dose-dependent role of Gfi1 in HSC function.

## Acknowledgements

The authors would like to thank Marina Suslo and Renata Köster for excellent technical assistance and the team of the animal facility at University Hospital Essen for genotyping, technical and administrative assistance during the whole mouse project. J.S. and the work were supported by a grant of the Fritz-Thyssen foundation. C.K. was supported by the IFORES fellowship of the University Hospital Essen, a Max-Eder fellowship from the German Cancer Fund (Deutsche Krebshilfe) as well as from the Dr. Werner Jackstädt-Stiftung.

## Author contribution

J.S. performed experiments and wrote the paper. A.T., Y.A.-M., P.K.P, D.F. performed experiments. D.R.E. und U.D. provided resources. C.K. conceived the study and supervised the project.

---

**Supplementary Figure 1:**
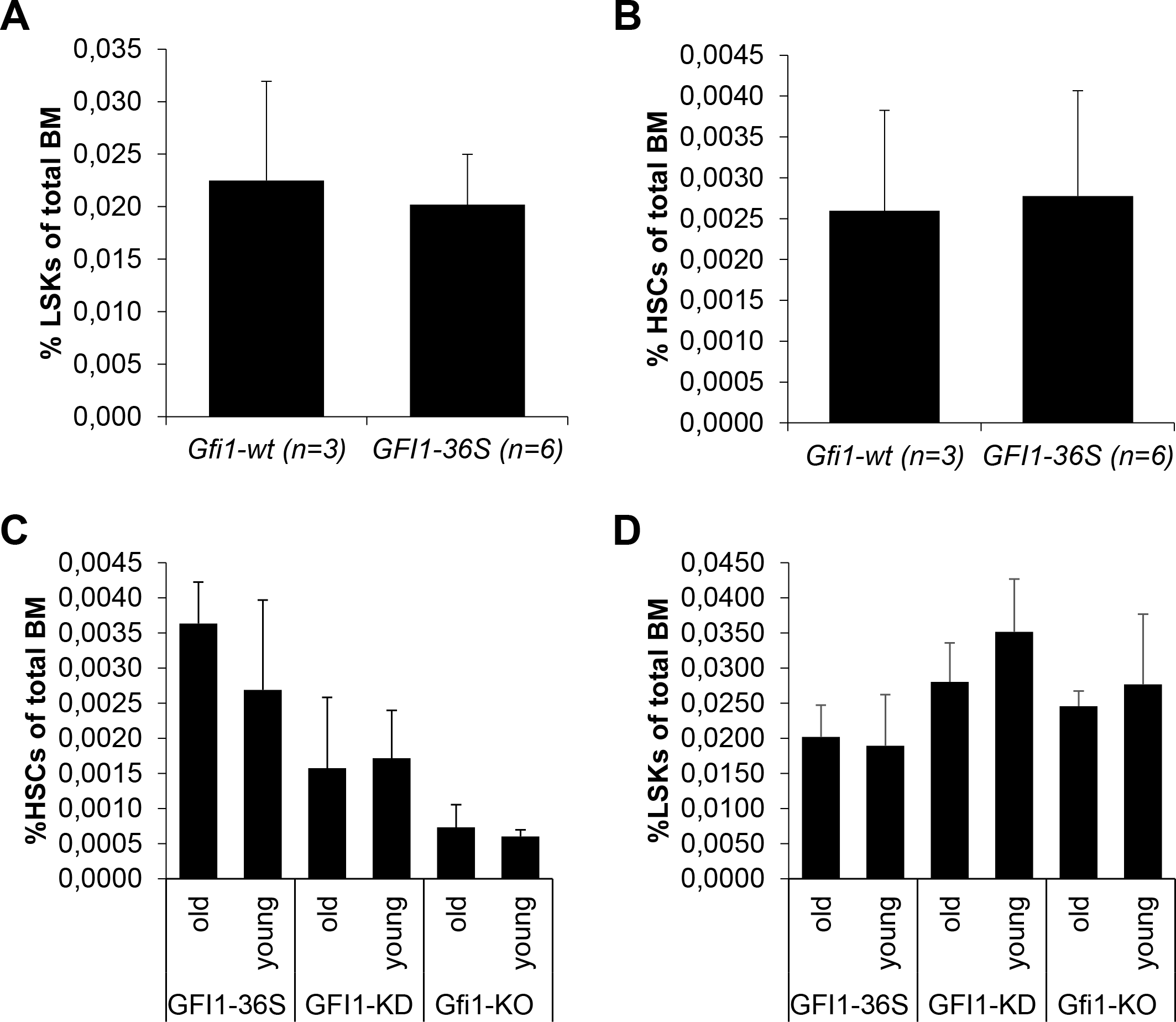
**A./B.** Number of LSK (Lin^−^c-Kit^+^Sca1^+^) cells (A) and HSCs (B) from *Gfi1*-wt mice containing the murine *Gfi1* gene are comparable with numbers of LSK cells or HSCs, respectively, from *GFI1*-36S mice in which the murine *Gfi1* gene was replaced by the human *GFI1* cDNA sequence. n = number of mice analyzed. No statistical difference detectable. **C./D.** The number of LSK cells (C) and HSCs (D) from *GFI1*-36S, *GFI1*-KD and *Gfi1*-KO mice did not change with age. Young = 39-127 days; old = 190-251 days. No statistical difference detectable.

**Supplementary Figure 2:**
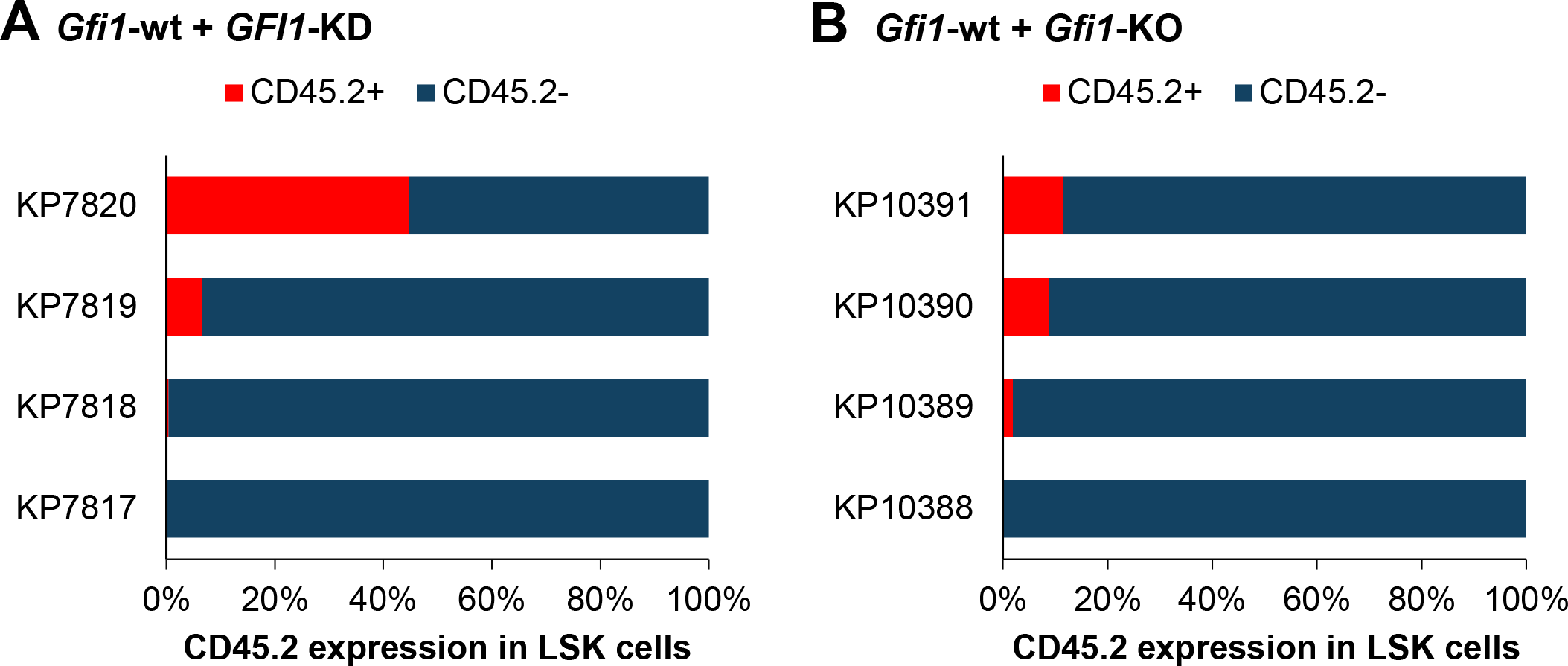
The differentiation defect is established early during hematopoiesis. **A.** CD45.2-expressing *GFI1*-KD cells rarely contributed to Lin^−^c-Kit^+^Sca1^+^ (LSK) cell compartment. Four mice with the indicated names (KP7817, KP7818, KP7819 and KP7820) were subjected to transplantation and BM cells were isolated 315 days (KP7819 and KP7920) or 322 days (KP7817 and KP7818) after transplantation. Cells were analyzed by flow cytometry. **B.** CD45.2-expressing *Gfi1*-KO cells hardly contribute to the generation of LSK cells. Four mice with the indicated names (KP10388, KP10389, KP10390 and KP10391) were subjected to transplantation and BM cells were isolated 112 days after transplantation. Cells were analyzed by flow cytometry.

